# Diagnosis of *Centrocestus formosanus* Infection in Zebrafish (*Danio rerio*): A Window on a New Globalization-Derived Invasive Parasite

**DOI:** 10.1101/814699

**Authors:** Antonino Pace, Ludovico Dipineto, Serena Aceto, Maria Concetta Censullo, Maria Carmen Valoroso, Lorena Varriale, Laura Rinaldi, Lucia Francesca Menna, Alessandro Fioretti, Luca Borrelli

**Affiliations:** Department of Veterinary Medicine and Animal Productions, Università degli Studi di Napoli Federico II, via Delpino 1, 80137, Naples, Italy; Department of Biology, Università degli Studi di Napoli Federico II, via Cintia 26, 80126, Naples, Italy

**Keywords:** Digenetic trematodes, Gill fluke, Invasive species, Molecular diagnosis, PCR, Zoonosis

## Abstract

*Centrocestus formosanus*is a digenetic trematode with a complex life cycle, involving invertebrate and vertebrate hosts, humans included. In particular, it causes gill lesions and mortality in freshwater fish species, and gastrointestinal symptoms in infected humans. Here,we describe the occurrence of *C. formosanus* infection in zebrafish imported in Italy. Gill arches of 30 zebrafish were examined for the presence of encysted metacercariae under a stereomicroscope, and processed through molecular analyses targeting the ribosomal Internal Transcribed Sequence 2 (ITS2) using species-specific primers of *C. formosanus*. Encysted metacercariae were found on the gills of 20/30 zebrafish and all were identified as *C. formosanus*.

Despite *C. formosanus* distribution was originally restricted to Asia, it has been subsequently reported in new countries, revealing itself as an invasive species, and raising important concerns for biodiversity, economy, animal and public health. Given the crucial role likely played by the ornamental fish industry in the spreading of this parasite, there is an urgent need for control measures to prevent the introduction and establishment of *C. formosanus* in non-endemic areas, Europe included. An adequate surveillance and health-monitoring program should be conducted in the development of microbiological and epidemiological approaches to diagnose and face these new globalization-derived invasive species.

## Introduction

Digenea are considered the largest group of internal metazoan parasites, including about 18,000 species [1]. They are almost ubiquitous, parasitizing a wide variety of vertebrate and invertebrate groups as intermediate or definitive hosts [1,2]. The importance of digenetic trematode infections in animals and humans has attracted much attention from different disciplines, in particular veterinary medicine. Indeed, most of digenetic trematode metacercariae are detected in freshwater fish hosts [2,3], and fish-borne zoonotic trematodes represent a concerning health issue in many Asian countries[2,4–6]. Already 20 years ago, the World Health Organization estimated more than 18 million of humans infected with fish-borne trematodes, and more than half a billion people at risk of infection worldwide[7,8]. In particular, the family Heterophyidae stood out for its clinical importance in humans, causing gastrointestinal and extra-intestinal infections[8–11].

Within fish-borne trematodes, *Centrocestus formosanus* is a small heterophyid fluke, described for the first time in Taiwan [12], and widely distributed in Asia [4,5,13–17]. Since the 1950’s,several authors have reported the introduction of this species in new continents, although its occurrence might still be underestimated [15,18–26].*C. formosanus*, similarly to other digenetic trematodes, exhibits a complex life cycle, as described by Nishigori[12]. The adults reside in the small intestine of vertebrate definitive hosts, such as birds and mammals. Eggs produced by adult trematodes hatch into miracidia, which use a thiarid snail as first intermediate host to develop into cercariae. Subsequently, free-swimming cercariae encyst in second intermediate fish hosts, specifically in the gills, where they develop into metacercariae. Piscivorous birds and mammals, ingesting the infected fish, complete the cycle[4,11-13,16,18,19,24–29]. Analogously, human infections might occur through consumption of raw or improperly cooked fish containing metacercariae[6].The parasite is highly specialized to its first intermediate snail host, *Melanoides tuberculata*[14,31]. On the contrary, several freshwater fish species might be infected, suggesting *C. formosanus* as a generalist parasite with a broad host range in second intermediate fish species[4,6,13,22–24,26,29,31,32].Therefore, pathogenicity evaluation is valuable for both wild and farmed(food and ornamental) fish[20]. Indeed, *C. formosanus*, causing severe lesions in the gills [23,24,26]with the resultant respiratory disorders, loss of production and death, is rightfully considered responsible for important economic losses in aquaculture [12,16,18,19,21–23,25,29,31-34].

Among the numerous freshwater fish species affected by *C. formosanus*[19,20,31], zebrafish (*Danio rerio*) has been suggested as susceptible to infection, but only three reports have been described to date [15,34,37], with a relative low prevalence (20%, 43%, and 5% respectively).

*Danio rerio* is a freshwater fish native to Asia, although it is widely distributed worldwide, probably due to aquarists’ and researchers’ predilection for it in home aquaria as well as animal model[38]. Indeed, the similarities between zebrafish and mammals led to a rapid increase in the use of zebrafish in scientific research, especially in developmental biology, neuroscience and behavioral research, and it proved to be more advantageous over previous model organisms[38,39].

The on-going growth of the ornamental fish industry, which includes more than 120 countries in the import and export of approximately 1.5 billion ornamental fish per year [40], has led to the appearance of problems in supply, traceability, sustainability, susceptibility to disease, and antibiotic resistance, which affect the trade[41].

Given the importance of *C. formosanus* infection and dissemination for animal and public health, and the implications for aquaculture, research and food safety, epidemiological investigations should be conducted in new geographical regions, in order to implement appropriate preventive and control measures [18].

The present study reports on the occurrence of *C. formosanus metacercariae* in the gills of zebrafish previously intended for research. To the authors’ knowledge, this is the second report of *C. formosanus* infections in zebrafish imported in Italy[37].Since the scarce and fragmentary data present in the European literature [29,42]are probably due to underestimated and under diagnosed expert evaluation, we propose to increase the awareness and ameliorate the diagnostic investigations to shed light on this zoonosis by morphological and molecular approach. In particular, we propose for the first time a fast and specific diagnostic method based of species-specific PCR primers to detect the presence of this new invasive species.

## Material and Methods

### Animal Maintenance

A total of 30 zebrafish, previously intended for research, was examined. All fish were male and female adult (4-6 month old) of heterozygous “wild type” strain, obtained from local commercial distributor. Fish were housed in groups of ten per 30 L tank, previously filled with deionized water, following an acclimation period of two weeks. Fish were fed two times daily with sterilized commercial food (Sera Vipagran, Germany). The room and water temperatures were maintained at 25–27 °C. Illumination (1010 ± 88 lx) was provided by ceiling-mounted fluorescent light tubes on a 14-h cycle (D:N = 14h:10h) consistent with the standards of zebrafish care[39,43].

Fish were treated in accordance with the Directive of the European Parliament and of the Council on the Protection of Animals Used for Scientific Purposes (directive 2010/63/EU) and in agreement with the Bioethical Committee of University Federico II of Naples(authorization protocol number 47339-2013).

During standard physical examination, performed under anesthesia by immersion in ethyl 3-aminobenzoate methane sulfonic acid solution (MS-222 at dose of 0.168 mg/ml) [39,43], the gills of 20 zebrafish were found to be affected by small white spots, ascribable to parasitic cysts (Fig. 1).Therefore, fish were not destined to research activities. The animals were euthanized by immersion in overdose 500 mg/ L of MS-222 buffered to pH 7.4 (Sigma– Aldrich, USA).

**Fig. 1.**
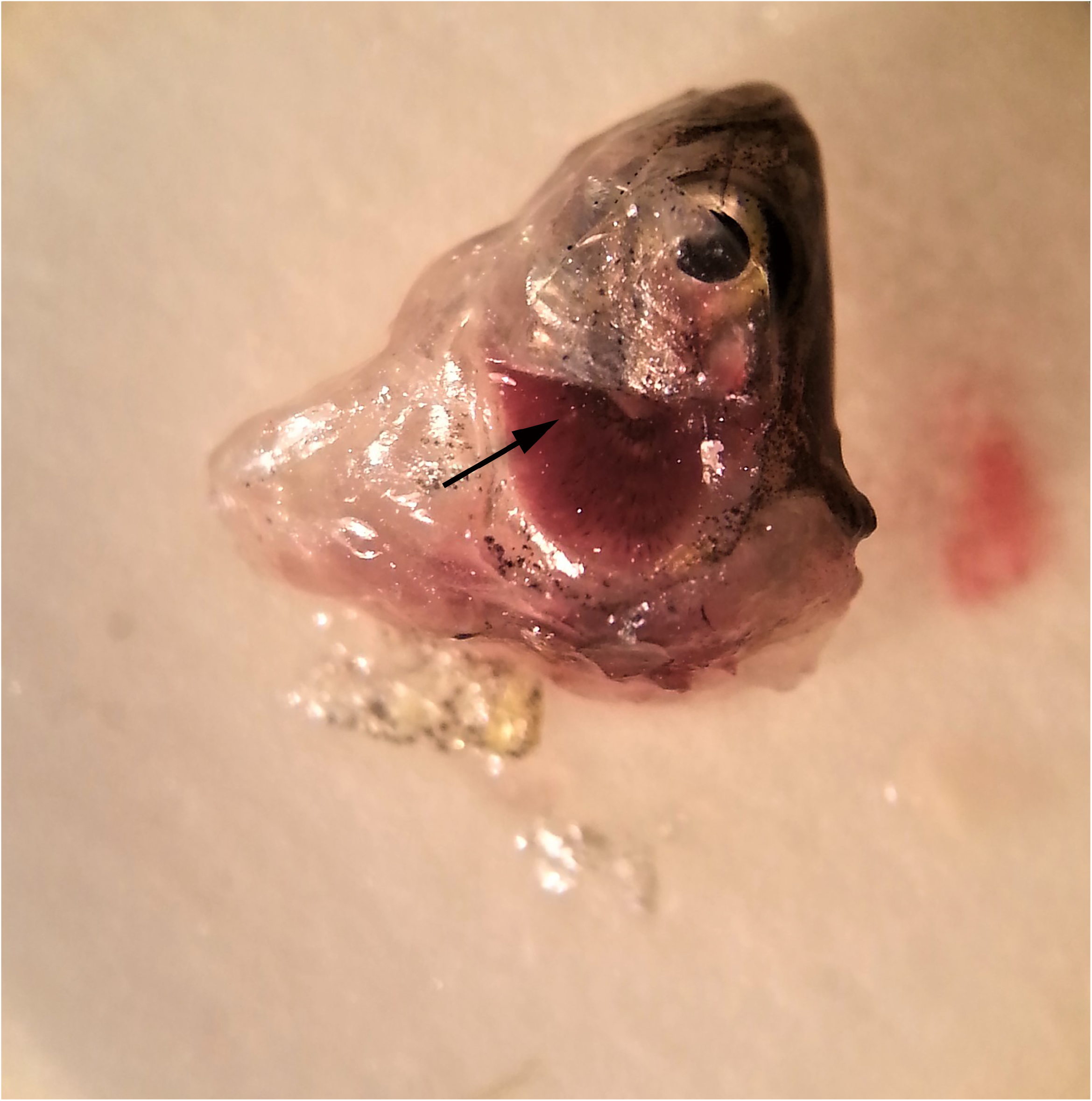
Gills of an infected zebrafish. Parasitic cysts are visible as miliar white lesions on the gill tissue(black arrow).

### Centrocestus formosanus Examination and Identification

Fish bodies were dissected, as reported in Borrelli et al. [39], and gill arches were removed with the aid of a stereomicroscope and prepared as wet mounts to be examined for the presence of encysted metacercariae[6,10,13,15,23,24,31,32,34,35]. Encysted metacercariae were examined under a light microscope to evaluate their morphology (Fig. 2)[10,44,45] and identified according to published characteristics [6,11-13–15,20,24,25,30] Live encysted metacercariae were also recorded at 40X using a Leica light microscope(S1 Video).

**Fig. 2.**
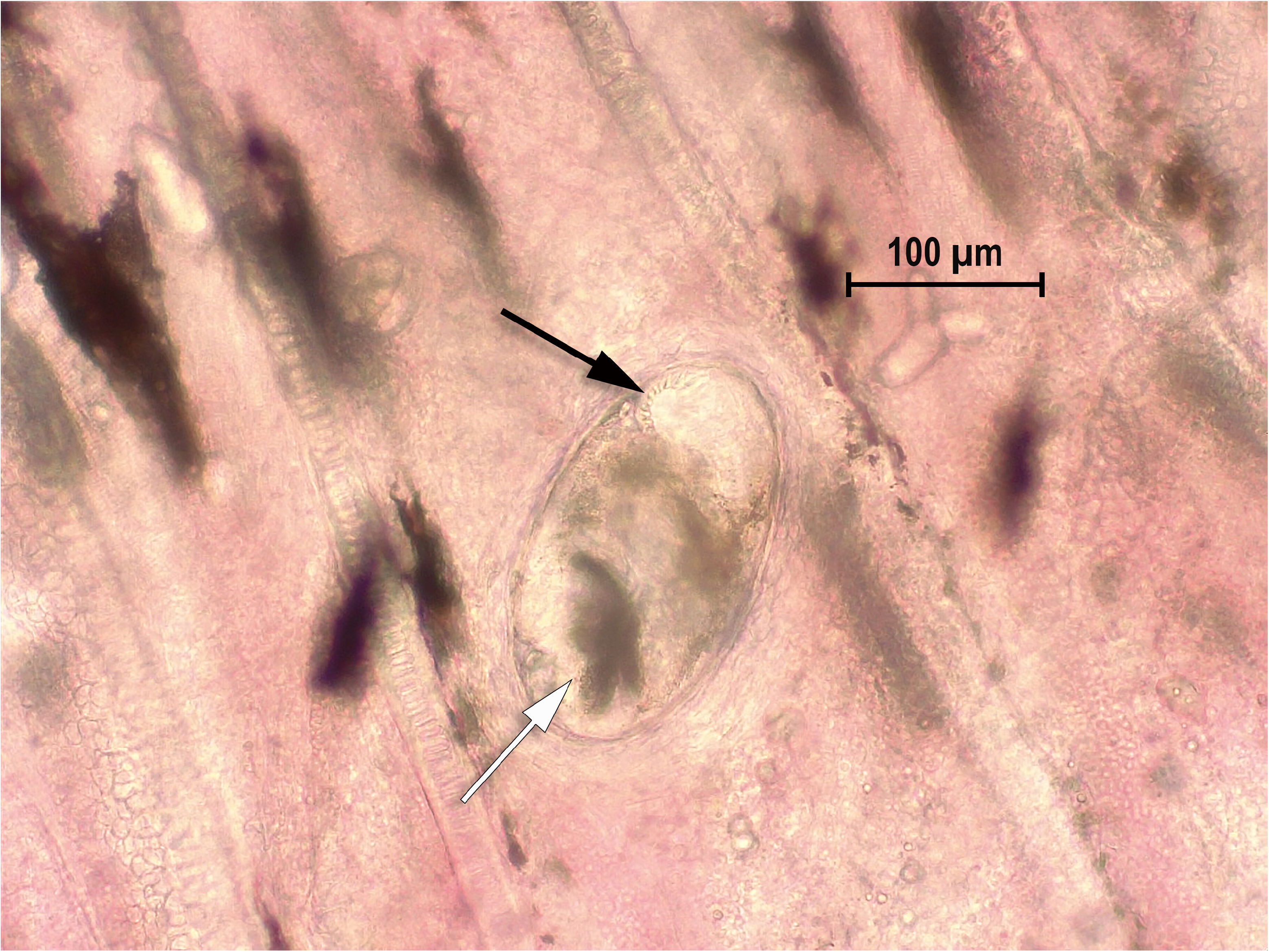
Encysted metacercaria in the gill tissue of infected zebrafish. 40X microscopy evaluation. Note the X-shaped excretory bladder (white arrow) and part of the oral sucker (black arrow).

### PCR Amplification and Sequencing

Total genomic DNA was extracted from 30 mg of gill tissue by using the QIAamp DNA Mini Kit (Qiagen). DNA concentrations and quality was assessed by spectrophotometric measurements with NanoDrop (ThermoFisher Scientific Inc.). DNA samples were stored at −20 °C until processed for amplification. The detection of *C. formosanus* DNA was performed by polymerase chain reaction (PCR) targeting the ribosomal Internal Transcribed Region 2 (ITS2), using the primer pair 3S (5’-GGTACCGGTGGATCACTCGGCTCGTG-3’) and BD2 (5’-TATGCTTAAATTCAGCGGGT-3’), previously described [15,37].As these primers are not specific for *C. formosanus*, we designed the species-specific primers ITS2_Centr_F 5’-ATGAAGAGCGCAGCCAACT-3’ and ITS2_Centr_R 5’-CGTGCAATGTTTGCATCGGA-3’to amplify a 393 bp fragment of the ITS2 region. PCR products were visualized by 1.5% agarose gel electrophoresis. Subsequently, the amplicons were cloned into the pSC-A-amp/kan vector (Agilent), sequenced using the T3 and T7 plasmid primers and analyzed using an ABI 310 Genetic Analyzer (Applied Biosystems). The obtained sequences were examined through BLAST analysis.

## Results

Over the total of 30 healthy zebrafish examined, 20 showed miliar gill cystic lesions (Fig. 1). Cysts were small and elliptical. Inside the cysts, the coiled mature metacercariae were characterized by a large, dark, X-shaped excretory bladder (occupying the majority of the body caudal portion) and by 32 circumoral spines surrounding the oral sucker, arranged in two rows. The shape of the excretory bladder and the number of circumoral spines are considered the most reliable characteristics in species identification within the genus *Centrocestus*, with *C. formosanus* characterized by an X-shaped excretory bladder and 32 circumoral spines, (S1 Video and Fig. 2)[15,20,25,44,45].Nevertheless, given that larval morphology might be confounding and that counting the number of circumoral spines might be challenging in some specimens [25], molecular analysis was conducted to confirm their taxonomic attribution. All the infected samples were positive to PCR amplification of the ribosomal ITS2 region fragment. However, the nucleotide sequence of the amplicons obtained with the primer pair 3S/BD2 [15,37] corresponds to the ITS2 fragment of the host *Danio rerio*. On the contrary, the species-specific primers designed in the present work amplify a fragment of 393 bp showing ~99% nucleotide identity with the homolog ITS2 sequences of *C. formosanus* present in GenBank(S2).

## Discussion

The occurrence of *C. formosanus* infection in zebrafish in Italy underlines the importance of focusing the attention on this invasive parasite, since this is the second case reported in the Italian peninsula [37]. Despite its origin and distribution was initially restricted to Asia[4,5,12–15,17,20,21,25], *C. formosanus* has been subsequently reported from Central and South America, Australia, and more recently Europe [13–15,18– 21,23,25,26,31,33,45,46].Actually, to the authors’ knowledge, this is the fourth case of *C. formosanus* infection in freshwater fish imported in Europe [29,37,42]and one of the most relevant in terms of number of infected animals, confirming the introduction of this parasite in the European area, as well as the possible underestimation of these infections.

The worldwide expansion of *C. formosanus* should be concerning for its ability to infect both ecologically and commercially valuable fish species [20,23,24,31].Indeed, *C. formosanus* metacercariae were detected on the gills of many freshwater fish species, with varying degrees of prevalence and severity [4–6,13,15,16,20,21,23,25,29,32,34,36,47].It has been hypothesized that different immunological responses across fish species might be responsible for mechanisms of resistance to parasite infection, although it has not been clearly determined [20,26,31,34].The lesions produced are very similar in all susceptible fish species [23,24,26], consisting in gill hyperemia and congestion, lamellar fusion and subsequent distortion of the architecture and reduction of the respiratory surface[20,21,23,24,32,33].These lesions eventually lead to low tolerance to hypoxia, respiratory difficulties and death [21,25,29,34]. This is particularly important under stressful environmental conditions(e.g. high temperatures, overcrowding and low water exchange), usually experienced in fish farms, where afflicted fish might exhibit decreased feeding rate, delayed development and mortality, resulting in the reduction of production and important economic losses[15,18–20,29,33–36].

Concerning the definitive hosts, both mammals and birds (e.g. canids, cats, rodents, anurans, pigeons, chickens, ducks, herons, cormorants, pelicans) have been reported to be suitable natural or experimental hosts[10,13,18–20,24,25].Nevertheless, the majority of literature addressed the qualitative aspects of infection in definitive hosts, and only recently quantitative aspects have been investigated[10,27].*C. formosanus* has been described causing lesions (e.g. fusion of *villi*, hyperplasia of crypts, epithelial damage) in the small intestine of experimentally infected herons and rats [21]. Similarly to fish hosts, the intensity of infection and the immune status of the host seemed related to the degrees of damage in definitive hosts [45].

In this regard, humans are also at risk of infection, and fish-borne trematodes have emerged as public health problems in Asia, especially in riverside areas, where the riparian populations are infected by consumption of raw and/or undercooked fish, containing infective *larvae* (i.e., metacercariae) [7,29,45].*C. formosanus* was not as prevalent as other fish-borne trematodes, but cases of human infection were reported in the Lao PDR, Vietnam, Thailand, Korea [12,13,15,19,20,25,34,41].Symptoms might vary from epigastric pain to indigestion, occasionally accompanied by diarrhea, although the relationship of these symptoms with *C. formosanus* infection was unclear, because the described patients were also infected by other trematodes [5,22,28,29].To date, no human cases have been described in the areas recently invaded by this parasite (e.g. US, Mexico, Brazil), nor in Europe [29,45].However, further investigation should be conducted, in order to keep a high level of attention on this issue[6,20,45].

The recent occurrence of *C. formosanus* in new hosts and countries suggests that its range is still expanding, possibly ending up into areas where it was not present, Europe included[19,20,29].The causes of this global spread are still subject of debate: some authors pointed at the dissemination of the first intermediate host, *Melanoides tuberculata*, whereas others hold responsible the movements of birds and freshwater fish, including the trade of ornamental species [13,18–20,26,27,29,31,32,34,35]. On one hand, the snail *M. tuberculate* has been deliberately introduced (for food and bio-control) or accidentally released (from aquaculture, aquarium trade or ballast water) in Central and South America, subsequently spreading also in the southern USA, all countries where *C. formosanus* infections have been reported[13,18,20,23,26,30,35,48].Similarly, a *M. tuberculata* population has been reported from Germany, making this area suitable for the completion of *C. formosanus* cycle and its establishment [29,49].On the other hand, the importation of ornamental freshwater fish from countries where *C. formosanus* is endemic likely plays a crucial role in the spreading of this parasite [28].Indeed, Asian countries are the major traders in the ornamental fish industry, exporting a wide range of species into Europe, which represents the largest global trade bloc, with the United Kingdom as the main importer and Italy at the 6^th^position[29,41,50]. In 2017, the estimated number of aquaria in Europe was 16,565,000, corresponding to approximately 300 million ornamental fish[51]. Of more than 2500 species involved in the ornamental fish industry, over 60% are of freshwater origin, and *D. rerio* is listed among the 30 species dominating the market. The trade largely relies on captive-bred fish, but significant numbers of specimens are also collected from the wild [41]. The top ten freshwater fish suppliers to Europe are: Singapore, Israel, Japan, Indonesia, Thailand, Sri Lanka, Colombia, China, Vietnam and Malaysia [41,50], most of them involved with *C. formosanus* infections[4–6,13–15,18,21,22,24,25,29,35,45].

Therefore, the main concerns are the spreading of this parasite in European freshwater habitats, due to intentional or accidental release of infected imported fish, and the resulting environmental, economic, and health implications[29,31].Primarily, some aspects of biology and epidemiology should be further explored in Europe[14,20], such as the presence and distribution of intermediate and definitive hosts, for the maintenance of the life cycle, or the prevalence of infection, particularly in second intermediate fish hosts[5,26,30,35].Another important element to consider is the likelihood of future cases of human infection, even if to date there have been no reports [8,24].Several strategies, to prevent the introduction and establishment of *C. formosanus* into non-endemic areas, have been proposed and tested during the last years [19,35]. Teams of experts in “one health” control should be the first actors involved in applying good management and efficient measures, especially during the intentional movement of animals, such as border inspection, accompanying health certification, quarantine measures, and, if necessary, treatment (prior to export or upon arrival) and disinfection procedures [29,33,46]. Additionally, adequate strategies should be applied in aquaculture facilities, including training of traders and farmers, regular examination of farmed fish, elimination of snail populations, avoidance of dispersal of farmed fish, and prevention of access to other species, especially birds and mammals[15,18,20,29,30,33,34].

The current report draws the attention on *C. formosanus* as an invasive parasite, as well as on other species that might be similarly introduced in Europe, underlining the need of epidemiological studies and appropriate preventive and control programs, in order to monitor their occurrence and prevent their negative consequences for economy, biodiversity, and animal and public health [14,20,29,30,33,46,52]. For these reasons, we propose a fast, cheap and specific PCR-based method to assess the infection of *C. formosanus* in zebrafish starting from small pieces of gill tissue of the host and avoiding elaborate collection of the metacercariae. The use of the parasite-specific primers eludes the frequent problem of the amplification of the host DNA and makes this method suitable also to detect this invasive parasite in other potential hosts. Specific recommendations concerning the diseases in ornamental fish should be strictly followed, as reported by Passantino et al. [53]. The mobilization of these animals, given the potential transmission of zoonoses from one country to others, should be better considered on the basis of good practices in diagnosing these potential pathogens. This control program might preserve animal and human international health, limiting the introduction and transfer of aquatic organism pathogens. We suggest also a proficient clinic approach developing new strategies in microbiology and epidemiology to better explore this new globalization-derived invasive species.

## Acknowledgements

This research received no specific grant from any funding agency in the public, commercial, or not-for-profit sectors. We would like to acknowledge and thank Dr. Adriana Petrovici and Laura Pacifico, DVM, Ph. D students for their precious support in fish gills microscopy image acquisition.

## Supplementary Materials

**S1 Video**

Live metacercaria encysted in the gill tissue of infected zebrafish, recorded at 40X using a Leica light microscope.

**S2**

ITS2 sequences of *C. formosanus*.

## References

[1] Olson PD, Cribb TH, Tkach VV, Bray RA, Littlewood DTJ. 2003. Phylogeny and classification of the Digenea (Platyhelminthes: Trematoda). Int J Parasitol 33:733–55. doi:10.1016/S0020-7519(03)00049-3.

[2] Sohn WM, Na BK, Cho SH, Lee SW, Choi SB, Seok WS. 2015. Trematode metacercariae in freshwater fish from water systems of hantangang and imjingang in Republic of Korea. Korean J Parasitol 53:289–98. doi:10.3347/kjp.2015.53.3.289.

[3] Choe S, Park H, Lee D, Kang Y, Jeon HK, Eom KS. 2018. Infections with digenean trematode metacercariae in two invasive alien fish, micropterus salmoides and lepomis macrochirus, in two rivers in chungcheongbuk-do, republic of Korea. Korean J Parasitol 56:509–13. doi:10.3347/kjp.2018.56.5.509.

[4] Chai JY, Sohn WM, Na BK, Yong TS, Eom KS, Yoon CH, Hoang EH, Jeoung HG, Socheat 2014. Zoonotic Trematode metacercariae in fish from Phnom Penh and Pursat, Cambodia. Korean J Parasitol 52:35–40. doi:10.3347/kjp.2014.52.1.35.

[5] Chai JY, Sohn WM, Na BK, Park JB, Jeoung HG, Hoang EH, Htoon TT, Tin HH. 2017. Zoonotic trematode metacercariae in fish from yangon, Myanmar and their adults recovered from experimental animals. Korean J Parasitol 55:631–41. doi:10.3347/kjp.2017.55.6.631.

[6] Krailas D, Veeravechsukij N, Chuanprasit C, Boonmekam D, Namchote S. 2016. Prevalence of fish-borne trematodes of the family Heterophyidae at Pasak Cholasid Reservoir, Thailand. Acta Trop 156:79–86. doi:10.1016/j.actatropica.2016.01.007.

[7] World Health Organization. Control of foodborne trematode infections, WHO Tech. Rep. Ser. No. 849; 1995. p. 1–157.

[8] Pulido-Murillo EA, Furtado LF V., Melo AL, Rabelo ÉML, Pinto HA. 2018. Fishborne zoonotic trematodes transmitted by Melanoides tuberculata snails, Peru. Emerg Infect Dis 24:606–8. doi:10.3201/eid2403.172056.

[9] Sohn WM, Na BK, Cho SH, Ju JW, Lee SW, Seok WS. 2018. Infections with zoonotic trematode metacercariae in yellowfin goby, Acanthogobius flavimanus, from coastal areas of Republic of Korea. Korean J Parasitol 56:259–65. doi:10.3347/kjp.2018.56.3.259.

[10] Mati VLT, Pinto HA, de Melo AL. 2013. Experimental infection of swiss and akr/j mice with centrocestus formosanus (trematoda: heterophyidae). Rev Inst Med Trop Sao Paulo 55:133–6. doi:10.1590/s0036-46652013000200013.

[11] Thaenkham U, Phuphisut O, Pakdee W, Homsuwan N, Sa-nguankiat S, Waikagul J, Nawa Y, Dung DT. 2011. Rapid and simple identification of human pathogenic heterophyid intestinal fluke metacercariae by PCR-RFLP. Parasitol Int 60:503–6. doi:10.1016/j.parint.2011.09.004.

[12] Nishigori M. 1924. On a new trematode Stamnosoma formosanum n. sp. and its life history. Taiwan Igakkai. Zasshi 1924;234:181–228.

[13] Salgado-Maldonado G, Rodriguez-Vargas MI, Campos-Perez JJ. Metacercariae of Centrocestus formosanus (Nishigori, 1924) (Trematoda) in Freshwater Fishes in Mexico and their Transmission by the Thiarid Snail Melanoides tuberculata. Stud Neotrop Fauna Environ 1995;30:245–50. doi:10.1080/01650529509360963.

[14] Yousif F, Ayoub M, Tadros M, El Bardicy S. 2016. The first record of Centrocestus formosanus (Nishigori, 1924) (Digenea: Heterophyidae) in Egypt. Exp Parasitol 168:56–61. doi:10.1016/j.exppara.2016.06.007.

[15] Wanlop A, Wongsawad C, Prattapong P, Wongsawad P, Chontananarth T, Chai JY. 2017. Prevalence of centrocestus formosanus metacercariae in ornamental fish from Chiang Mai, Thailand, with molecular approach using ITS2. Korean J Parasitol 55:445–9. doi:10.3347/kjp.2017.55.4.445.

[16] Chai JY, Sohn WM, Jung BK, Yong TS, Eom KS, Min DY, Insisengmay B, Insisiengmay S, Phommasack B, Rim HJ. 2015. Intestinal helminths recovered from humans in Xieng Khouang Province, Lao PDR with a particular note on Haplorchis pumilio infection. Korean J Parasitol 2015; 53:439–45. doi:10.3347/kjp.2015.53.4.439.

[17] Komatsu S, Kimura D, Paller VGV., Uga S. 2014. Dynamics of Centrocestus armatus Transmission in Endemic River in Hyogo Prefecture, Japan. Trop Med Health 42:35–42. doi:10.2149/tmh.2013-34.

[18] Ximenes RF, Gonçalves ICB, Miyahira IC, Pinto HA, Melo AL, Santos SB. 2016.. Centrocestus formosanus (Trematoda: Heterophyidae) in Melanoides tuberculata (Gastropoda: Thiaridae) from Vila do Abraão, Ilha Grande, Rio de Janeiro, Brazil. Brazilian J Biol 77:318–22. doi:10.1590/1519-6984.13615.

[19] Pinto HA, Gonçalves NQ, López-Hernandez D, Pulido-Murillo EA, Melo AL. 2018. The life cycle of a zoonotic parasite reassessed: Experimental infection of Melanoides tuberculata (Mollusca: Thiaridae) with Centrocestus formosanus (Trematoda: Heterophyidae). PLoS One 13:1–13. doi:10.1371/journal.pone.0194161.

[20] Scholz T, Salgado-Maldonado G. 2000. The Introduction and Dispersal of Centrocestus formosanus (Nishigori, 1924) (Digenea: Heterophyidae) in Mexico: A Review. Am Midl Nat 143:185–200. doi:10.1674/0003-0031(2000)143[0185:tiadoc]2.0.co;2.

[21] Sumuduni BGD, Munasinghe DHN, Arulkanthan A. 2018. Chronological analysis of the damages caused by the metacercariae of Centrocestus formosanus in the gills of Cyprinus carpio and lesions caused by the adult flukes in Ardeola ralloides: An experimental study. Int J Vet Sci Med 6:165–71. doi:10.1016/j.ijvsm.2018.08.006.

[22] Chai JY, Yong TS, Eom KS, Min DY, Jeon HK, Kim TY, Jung BK, Sisabath L, Insisiengmay B, Phommasack B, Rim HJ. 2013. Hyperendemicity of Haplorchis taichui infection among riparian people in Saravane and Champasak province, Lao PDR. Korean J Parasitol 51:305–11. doi:10.3347/kjp.2013.51.3.305.

[23] Mitchell AJ, Salmon MJ, Huffman DG, Goodwin AE, Brandt TM. 2000. Prevalence and pathogenicity of a heterophyid trematode infecting the gills of an endangered fish, the fountain darter, in two central Texas spring-fed rivers. J Aquat Anim Health 12:283–9. doi:10.1577/1548-8667(2000)012<0283:PAPOAH>2.0.CO;2.

[24] Vélez-Hernández EM, Constantino-Casas F, García-Márquez LJ, Osorio-Sarabia D. 1998. Gill lesions in common carp, Cyprinus carpio L., in Mexico due to the metacercariae of Centrocestus formosanus. J Fish Dis 21:229–32. doi:10.1046/j.1365-2761.1998.00091.x.

[25] Wongsawad C, Wongsawad P, Sukontason K, Maneepitaksanti W, Nantarat N. 2017. Molecular phylogenetics of centrocestus formosanus (Digenea: Heterophyidae) originated from freshwater fish from chiang Mai Province, Thailand. Korean J Parasitol 55:31–7. doi:10.3347/kjp.2017.55.1.31.

[26] Mitchell AJ, Goodwin AE, Salmon MJ, Brandt TM. 2002. Experimental Infection of an Exotic Heterophyid Trematode, Centrocestus formosanus, in Four Aquaculture Fishes. N Am J Aquac 64:55–9. doi:10.1577/1548-8454(2002)064<0055:EIOAEH>2.0.CO;2.

[27] Pinto HA, Mati VLT, de Melo AL. 2015. Experimental centrocestiasis: Worm burden, morphology and fecundity of Centrocestus formosanus (Trematoda: Heterophyidae) in dexamethasone immunosuppressed mice. Parasitol Int 64:236–9. doi:10.1016/j.parint.2015.02.002.

[28] El-Azazy OME, Abdou NEMI, Khalil AI, Al-Batel MK, Majeed QAH, Henedi AAR. Tahrani LMA. 2015. Potential zoonotic trematodes recovered in stray cats from Kuwait municipality, Kuwait. Korean J Parasitol 53:279–87. doi:10.3347/kjp.2015.53.3.279.

[29] Mehrdana F, Jensen HM, Kania PW, Buchmann K. 2014. Import of exotic and zoonotic trematodes (Heterophyidae: Centrocestus sp.) in Xiphophorus maculatus: Implications for ornamental fish import control in Europe. Acta Parasitol 59:276–83. doi:10.2478/s11686-014-0237-z.

[30] Tolley-Jordan LR, Chadwick MA. 2019. Effects of Parasite Infection and Host Body Size on Habitat Associations of Invasive Aquatic Snails: Implications for Environmental Monitoring. J Aquat Anim Health 31:121–8. doi:10.1002/aah.10059.

[31] Frankel VM, Hendry AP, Rolshausen G, Torchin ME. 2015. Host preference of an introduced “generalist” parasite for a non-native host. Int J Parasitol 45:703–9. doi:10.1016/j.ijpara.2015.03.012.

[32] Huston DC, Cantu V, Huffman DG. 2014. Experimental Exposure of Adult San Marcos Salamanders and Larval Leopard Frogs to the Cercariae of Centrocestus formosanus. J Parasitol 100:239–41. doi:10.1645/13-419.1.

[33] Soler-Jiménez LC, Paredes-Trujillo AI, Vidal-Martínez VM. 2017. Helminth parasites of finfish commercial aquaculture in Latin America. J Helminthol 91:110–36. doi:10.1017/s0022149x16000833.

[34] Ortega C, Fajardo R, Enríquez R. 2009. Trematode centrocestus formosanus infection and distribution in ornamental fishes in Mexico. J Aquat Anim Health 21:18–22. doi:10.1577/H07-022.1.

[35] Pinto HA, Mati VLT, Melo AL. 2014. Metacercarial infection of wild nile tilapia (Oreochromis niloticus) from Brazil. Sci World J. doi:10.1155/2014/807492.

[36] Mendoza-Estrada LJ, Hernández-Velázquez VM, Arenas-Sosa I, Flores-Pérez FI, Morales-Montor J, Penã-Chora G. 2016. Anthelmintic Effect of Bacillus thuringiensis Strains against the Gill Fish Trematode Centrocestus formosanus. Biomed Res Int. doi:10.1155/2016/8272407.

[37] Iaria C, Migliore S, Macri D, Bivona M, Capparucci F, Gaglio G, Marino F. 2019. Evidence of Centrocestus formosanus (Nishigori, 1924) in Zebrafish (Danio rerio). Zebrafish. doi:10.1089/zeb.2019.1744.

[38] Kinth P, Mahesh G, Panwar Y. 2013. Mapping of Zebrafish Research: A Global Outlook. Zebrafish 10:510–7. doi:10.1089/zeb.2012.0854.

[39] Borrelli L, Aceto S, Agnisola C, De Paolo S, Dipineto L, Stilling RM, Dinan TG, Cryan JF, Menna LF, Fioretti A. 2016. Probiotic modulation of the microbiota-gut-brain axis and behaviour in zebrafish. Sci Rep 6:1–9. doi:10.1038/srep30046.

[40] Ornamental Fish International. Retrieved from: https://www.ofish.org/ornamental-fish-industry-data.

[41] Dey VK. 2016. The Global Trade in Ornamental Fish. Infofish 4:52–5.

[42] Gjurčević E, Petrinec Z, Kozarić Z, Kuzir S, GjurčevićK, Vičemilo M, Dzaja P. 2007. Metacercariae of Centrocestus formosanus in goldfish (Carassius auratus L.) imported into Croatia. Helminthologia 44:214–6. doi:10.2478/s11687-007-0034-4.

[43] Westerfield M. 2007. The Zebrafish Book: A Guide for the Laboratory Use of Zebrafish (*Danio rerio*). Eugene (OR): University of Oregon Press.

[44] Sohn WM, Na BK, Cho SH, Ju JW, Kim CH, Yoon KB, Kim JD, Son DC, Lee SW. 2018. Infections with Centrocestus armatus metacercariae in fishes from water systems of major rivers in republic of Korea. Korean J Parasitol 56:341–9. doi:10.3347/kjp.2018.56.4.341.

[45] Chai JY, Sohn WM, Yong TS,Eom KS, Min DY, Lee MY, Lim H, Insisiengmay B, Phommasack B, Rim HJ. 2013. Centrocestus formosanus (Heterophyidae): Human Infections and the Infection Source in Lao PDR. J Parasitol 99:531–6. doi:10.1645/12-37.1.

[46] Evans BB, Lester RJG. 2001. Parasites of ornamental fish imported into Australia. Bull Eur Assoc Fish Pathol 21:51–5.

[47] Eom KS, Park HS, Lee D, Sohn WM, Yong TS, Chai JY, Min DY, Rim HJ, Insisiengmay B, Phommasack B. 2015. Infection status of zoonotic trematode metacercariae in fishes from Vientiane municipality and Champasak Province in Lao PDR. Korean J Parasitol 53:447–53. doi:10.3347/kjp.2015.53.4.447.

[48] Clusa L, Miralles L, Basanta A, Escot C, García-Vázquez E. 2017. eDNA for detection of five highly invasive molluscs. A case study in urban rivers from the Iberian Peninsula. PLoS One 12:1–14. doi:10.1371/journal.pone.0188126.

[49] Glöer P 2002. Die Süsswassergastropoden Nord-und Mitteleuropas: Bestimmungsschlüssel, Lebensweise, Verbreitung. Die Tier welt Deutschlands. 73. Teil. Conch Books, Hackenheim, Germany, 327 pp.

[50] OATA. 2017. EU Ornamental Fish Import & Export Statistics 2016 (Third Countries & Intra-EU Community trade). Westbury (UK).

[51] FEDIAF. 2017. European Facts&Figures2017. Bruxelles.

[52] Chai JY, Murrell KD, Lymbery AJ. 2005. Fish-borne parasitic zoonoses: Status and issues. Int J Parasitol 35:1233–54. doi:10.1016/j.ijpara.2005.07.013.

[53] Passantino A, Macrì D, Coluccio P, Foti F, Marino F. 2008. Importation of mycobacteriosis with ornamental fish: Medico-legal implications. Travel Med Infect Dis 6:240–4. doi:10.1016/j.tmaid.2007.12.003.

